# Molecular connectivity between extra-cytoplasmic sigma factors and PhoP accounts for integrated mycobacterial stress response

**DOI:** 10.1101/2021.10.25.465832

**Authors:** Harsh Goar, Partha Paul, Hina Khan, Dibyendu Sarkar

## Abstract

The main purpose of this study is to understand how mycobacteria can sense numerous stress conditions and mount an appropriate stress response. Recent studies suggest that at low pH *M. tuberculosis* encounters reductive stress, and in response, modulates redox homeostasis by utilizing the *phoPR* regulatory system. However, the mechanism of integrated regulation of stress response remains unknown. To probe how PhoP contributes to redox stress response, we find that a PhoP-depleted *M. tuberculosis* shows a significantly enhanced susceptibility to redox stress relative to the WT bacilli. In keeping with these results, PhoP was shown to contribute to mycothiol redox state. Because SigH, one of the alternative sigma factors of mycobacteria, is known to control expression of redox inducible genes, we probed whether previously-reported PhoP-SigH interaction accounts for mycobacterial redox stress response. We had shown that under acidic conditions PhoP functions in maintaining pH homeostasis via its interaction with SigE. In striking contrast, here we show that under redox stress, direct recruitment of SigH, but not PhoP-SigH interaction, controls expression of mycobacterial thioredoxin genes, a major mycobacterial anti-oxidant system. Together, these unexpected results uncover novel stress-specific enhanced or reduced interaction events of sigma factors and PhoP, as the underlying mechanisms of an adaptive programme, which couples low pH conditions and mycobacterial thiol redox homeostasis.

**Significance:** *M. tuberculosis* encounters reductive stress under acidic pH. To investigate the mechanism of integrated stress response, we show that PhoP plays a major role in mycobacterial redox stress response. We observed a significant correlation between *phoP*-dependent and redox-active expression of thioredoxin genes, a major mycobacterial antioxidant system. Further probing on functioning of regulators reveals that while PhoP controls pH homeostasis via its interaction with SigE, direct recruitment of SigH, but not PhoP-SigH interaction, controls expression of thioredoxin genes. These strikingly contrasting results showing enhanced PhoP-SigE interaction under acidic pH and reduced PhoP-SigH interaction under redox conditions, uncover the underlying novel mechanism of mycobacterial adaptive program, coupling low pH with maintenance of redox homeostasis.

## Introduction

Survival and persistence of mycobacteria under varying environmental cues rely on the coupling between signal sensing and induction of appropriate adaptive program. Therefore, understanding molecular mechanism of adaptation in response to environmental changes represents a major aspect of TB biology. Although recent years have seen substantial progress in the molecular characterization of the bacillus, much work is still needed to understand how *M. tuberculosis* copes with varying environments it encounters in the course of an infection. Adaptations to such conditions must require a complex regulatory mechanism of gene expression.

Endogenous oxidative stress represents a significant challenge against survival and growth of microbes adapted to an aerobic life style (1). However, intracellular pathogens like *M. tuberculosis*, also encounters exogenous oxidative stress generated by the host (2). Expectedly, *M. tuberculosis* is equipped with a number of dedicated anti-oxidant systems to ensure survival and growth within the host macrophages (3-10). Among these, the thioredoxin system coupled with glutathione controls many important cellular processes, such as antioxidant pathways, DNA and protein repair enzymes, and activation of redox-sensitive transcription factors (11, 12). Although many Gram negative bacteria possess both systems, the glutathione system is lacking in *M. tuberculosis* (7, 12), and mycothiol is considered as substitute for glutathione (7). While mycothiol-deficient *M. tuberculosis* is not significantly growth attenuated in mice (13), thioredoxin reductase (TrxB2) is essential for *in vitro* growth, underscoring the importance of TrxB2 (14-16). Consistent with these results, under physiological conditions, depletion of TrxB2 causes lytic death of mycobacteria and is essential for growth and survival under *in vitro* and in mice (17). However, we still do not understand the regulatory mechanism of control of thioredoxin system under redox stress.

Mycobacterial gene expression in response to acidic pH significantly overlaps with the regulon, which is under the control of PhoPR two-component system (18). In keeping with this, a large subset of low pH-inducible PhoPR-regulated genes are induced immediately following *M. tuberculosis* phagocytosis and remain induced during macrophage infection (19-22). These include genes involved in biosynthesis of cell envelope lipids, which contribute to arresting phagosomal maturation (23), indicating a major shift in anabolic metabolism under acidic pH. PhoPR, in addition, controls major virulence factors central to TB pathogenesis, including ESX-1 dependent secretion of ESAT-6 (24-26), which interferes with phagosomal maturation, neutralizes phagosome and modulates carbon source availability in the mycobacterial cytoplasm from the fatty acids and cholesterol of the phagosome (27). Thus, during onset of macrophage infection, PhoPR activation is linked to acidic pH, and available carbon source, suggesting a physiological link between pH, carbon source and macrophage pathogenesis.

Given the relationship that exists between PhoP and SigH as the regulators of mycobacterial redox response on one hand, and integration of pH stress and reductive stress to govern metabolic plasticity of mycobacteria on the other hand, we sought to investigate role of the *phoP* locus in mycobacterial redox stress response. We found a significant correlation between *phoP*-dependent and redox-specific expression of mycobacterial thioredoxin genes. In agreement with this, a PhoP-depleted mutant is significantly more susceptible to redox stress relative to the WT bacilli. Based on further probing on functioning of regulators, we suggest a model which provides new biological insights into the metabolic events necessary to maintain mycobacterial thiol redox homeostasis. Although PhoP controls pH homeostasis via its interaction with SigE (28), here we demonstrate that direct recruitment of SigH alone, but not PhoP-SigH interaction, regulates expression of mycobacterial thioredoxin genes. Taken together, these results uncover the underlying mechanism of an adaptive programme, which couples mycobacterial thiol redox homeostasis with low pH conditions.

## RESULTS

### Modulation of mycobacterial redox environment is controlled by PhoP

Recent studies demonstrate that under low pH conditions induction of *phoPR* triggers restricted metabolism utilizing specific carbon source, resulting a slower bacterial growth, and modulation of redox homeostasis to counter reductive stress associated with acidic pH (27, 29). Thus, we investigated the impact of PhoP depletion on susceptibility of *M. tuberculosis* to redox stress. To this end, WT-H37Rv, Δ *phoP-*H37Rv and the complemented mutant strains were grown in presence of increasing concentrations of diamide, a thiol-specific oxidant and mycobacterial survival was monitored by microplate-based assays using Alamar Blue, an oxidation-reduction indicator (SI Appendix, Fig. S1A). In this assay, change of a non-fluorescent blue to a fluorescent pink appearance is directly linked to bacterial growth. We next quantified and plotted the fluorescence data to determine fold difference in susceptibility of WT and the mutant bacilli to diamide (SI Appendix, Fig. S1B). Note that bacterial survival under normal conditions of growth (in presence of carry-over concentrations of DMSO) and under redox stress (in presence of 5 mM diamide) were assessed by normalizing the data with respect to corresponding WT-H37Rv cultures (considered as 100%). For *ΔphoP*-H37Rv, a significantly lower growth (< 10%) relative to the WT bacilli in presence of diamide suggests role of PhoP as a major regulator of mycobacterial redox stress response. Importantly, the growth defect of the mutant under redox stress could be restored by stable expression of a copy of the *phoP* gene (as described in the SI Appendix, SI Methods), and the mutant grew comparably well as that of the WT bacteria under normal conditions of growth. Together, these results suggest a specific role of PhoP in detoxifying thiol-oxidizing stress.

To further validate our results, we grew WT and PhoP-depleted *M. tuberculosis* H37Rv (*ΔphoP*-H37Rv) in 5 mM diamide for 48 hours and measured their survival by enumerating CFU values (Fig. 1A). The results highlight that in the presence of diamide, *ΔphoP*-H37Rv displayed a significantly stronger growth inhibition (4.5±2 - fold) relative to WT-H37Rv. More importantly, stable PhoP expression could significantly restore diamide-dependent growth inhibition of *ΔphoP*-H37Rv, suggesting that PhoP plays a major role in detoxifying redox stress. Likewise, in agreement with a previously-reported result (20), we noted a significantly enhanced sensitivity of Δ *phoP*-H37Rv to 50 μM Cumene Hydrogen Peroxide (CHP) relative to the WT-H37Rv and complementation of *ΔphoP*-H37Rv completely rescued sensitivity of the mutant bacilli to CHP (Fig. 1B).

**Fig. 1:**
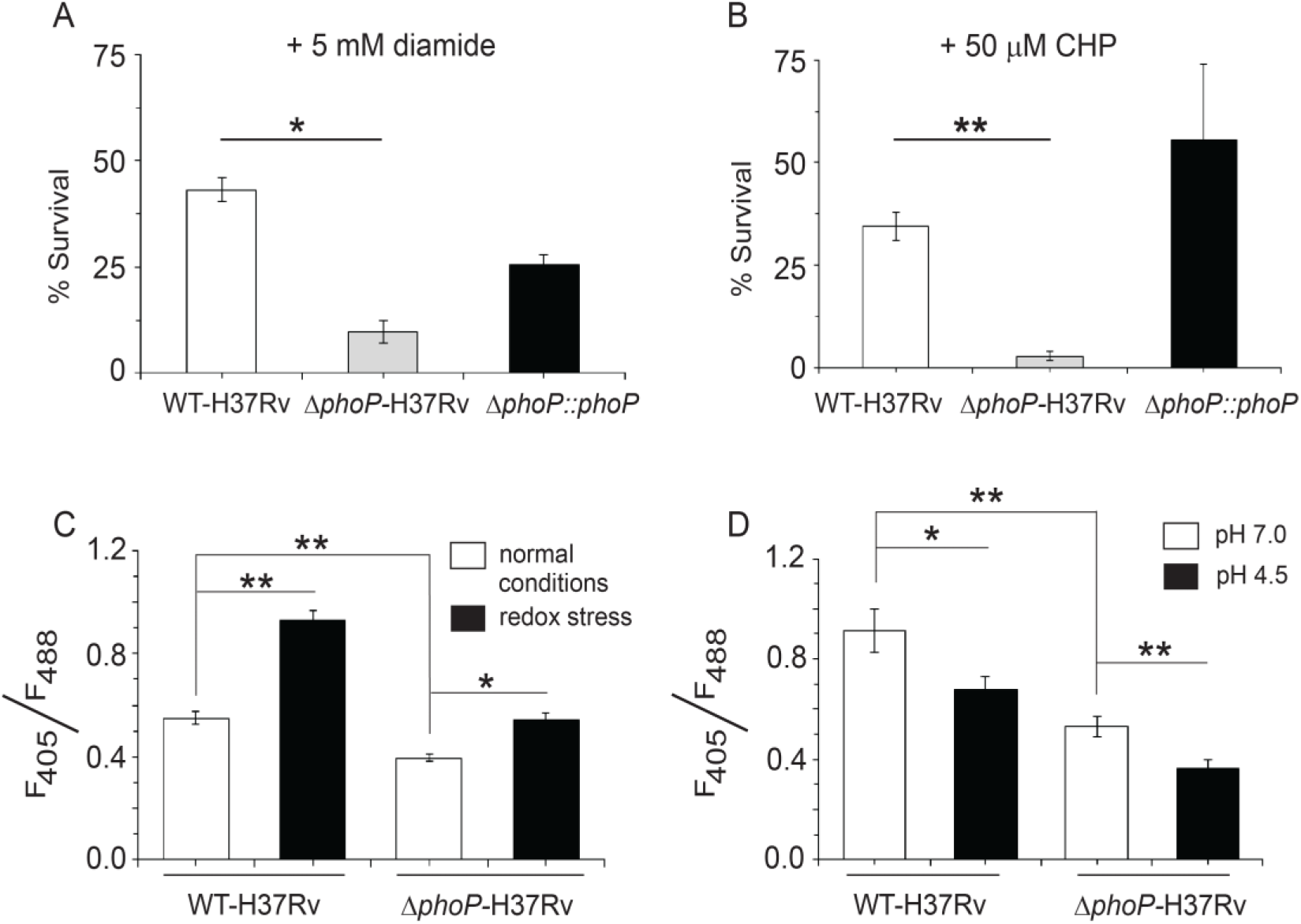
*phoP* plays a major role in mycobacterial redox stress response. (A-B) To compare susceptibility to stress conditions, WT-H37Rv and Δ *phoP*-H37Rv were grown either in presence of 5 mM Diamide for 48 hours or 50 µM Cumene hydroperoxide (CHP) for 24 hours, and CFU values were enumerated. Note that for each strain percent bacterial survival was determined relative to growth of the corresponding strain in presence of carryover DMSO. The results suggest that Δ *phoP* is significantly growth defective (relative to WT bacteria) both in presence of diamide and CHP; however, the growth defect of the mutant is largely rescued in the Δ *phoP-*complemented strain, suggesting significantly enhanced susceptibility of the mutant bacilli under redox stress. The growth experiments were performed in biological duplicates, each with two technical repeats. To examine whether *phoP* is linked to mycobacterial redox potential, we compared intra-mycobacterial mycothiol redox state of WT and Δ *phoP*-H37Rv grown either under (C) redox stress or (D) low pH. In this experiment, we used plasmid Mrx1-roGPF2 where mycoredoxin is fused to redox-sensitive GFP enabling real time measurement of mycothiol redox state in mycobacteria (34). The biosensor response was measured by analysis of fluorescence emission at 510 nm after excitation of the samples at 405 nm and 488 nm, and the results show average values from biological triplicates, each with two technical repeats (^*^P≤0.05; ^**^P≤0.01). The results strongly suggest that mycobacterial *phoP* locus contributes to mycothiol redox potential.

Next, we grew WT-H37Rv and *ΔphoP*-H37Rv in presence of 5 mM Diamide to investigate redox-dependent change in mycothiol redox state (Fig. 1C). While most eukaryotes and a large number of prokaryotes utilize glutathione (GSH) to maintain redox balance, genus Mycobacterium produces two different low molecular weight thiols i.e., mycothiol (MSH) and ergothioneine (ERG). The non-protein thiol MSH, which is produced in millimolar concentration in cells (30), is capable of reducing oxidized cysteine residues of proteins by mycoredoxin (30-33). In keeping with this, MSH mutants are more susceptible to nitrosative and redox stress compared to WT bacilli. Importantly, development of a green fluorescence-based sensor (GFP) where mycoredoxin is fused to redox-sensitive GFP, has enabled real-time measurement of mycothiol redox state in mycobacteria (34).

Singh and co-workers have utilized this probe to demonstrate that acidic pH inside phagosomes, induces reductive stress in *M. tuberculosis* residing within macrophages (35). Using this probe, our results demonstrate that mycothiol redox state of WT-H37Rv remains significantly higher relative to Δ *phoP*-H37Rv grown under normal conditions, suggesting that PhoP contributes to mycobacterial thiol redox homeostasis (Fig. 1C). Notably, WT-H37Rv shows a significant increase in mycothiol redox state when grown in presence of 5 mM diamide relative to cells grown under normal conditions (1.7±0.02-fold) (Fig. 1C). In contrast, *ΔphoP*-H37Rv shows reduced levels (1.3±0.03-fold) of mycothiol redox state for cells grown in presence of 5 mM diamide relative to normal conditions (Fig. 1C). We consider this difference significant as minor change in ratios of fluorescence intensities often impact intracellular redox state of mycobacteria (35-38). Because *ΔphoP*-H37Rv displayed a significantly lower intra-mycobacterial mycothiol redox state relative to WT-H37Rv under normal conditions as well as under redox stress, we conclude that most likely PhoP plays a role in mycobacterial thiol redox homeostasis.

Having shown a striking effect of *phoP* locus on mycobacterial susceptibility to redox stress, we next measured intra-mycobacterial mycothiol redox state of WT- and *ΔphoP*-H37Rv grown under low pH conditions (Fig. 1D). For the WT bacilli, we observed a significantly (1.3±0.1 -fold) lower mycothiol redox state under low pH conditions, relative to the bacilli grown under normal conditions (pH 7.0). This is consistent with mycobacterial adaptation under low pH conditions which lead to reductive stress (35, 39). In contrast, *ΔphoP*-H37Rv under identical conditions showed 1.5±0.04-fold lower mycothiol redox state compared to the mutant grown under normal pH (pH 7.0), suggesting that low pH-induced alteration in redox environment requires the presence of *phoP* locus to maintain mycobacterial thiol redox homeostasis.

### Mycobacterial survival under redox stress requires SigH

To investigate diamide-induced global gene expression profile, WT-H37Rv was grown to mid log phase (0.4 – 0.5 A_600_) and then treated with 5 mM diamide or an equivalent volume of DMSO. After 2 hours of growth, total RNA was isolated and subjected to RNA sequencing. Our results identified 25 significantly upregulated (>3.5-fold; p<0.01) genes (Fig. 2A) [SI Appendix, Table S3A (Excel spreadsheet)]. Most of these belong to either heat stress response (*hsp, Rv2466c, Rv3054c*), or oxidative stress response (*trxB1, trxB2, sigE*, and *SigH*). Because sigma factors are central to much of the success of *M. tuberculosis* to adapt to varying host environments through complex transcriptional programs, we compared global expression profile of sigma factors for bacterial cells grown with or without redox stress (Fig. 2B). Consistent with RNA-seq data (Fig. 2A), two sigma factors (SigE and SigH) out of a total of 13 showed approximately ∼30-50-fold redox-inducible activation of expression. Although SigH expression showed the highest level of redox-dependent activation, under identical conditions other sigma factor encoding genes did not display a significant difference in their expression. These observations are consistent with previous studies suggesting a major role of SigH in mycobacterial redox stress response (40, 41).

**Fig. 2:**
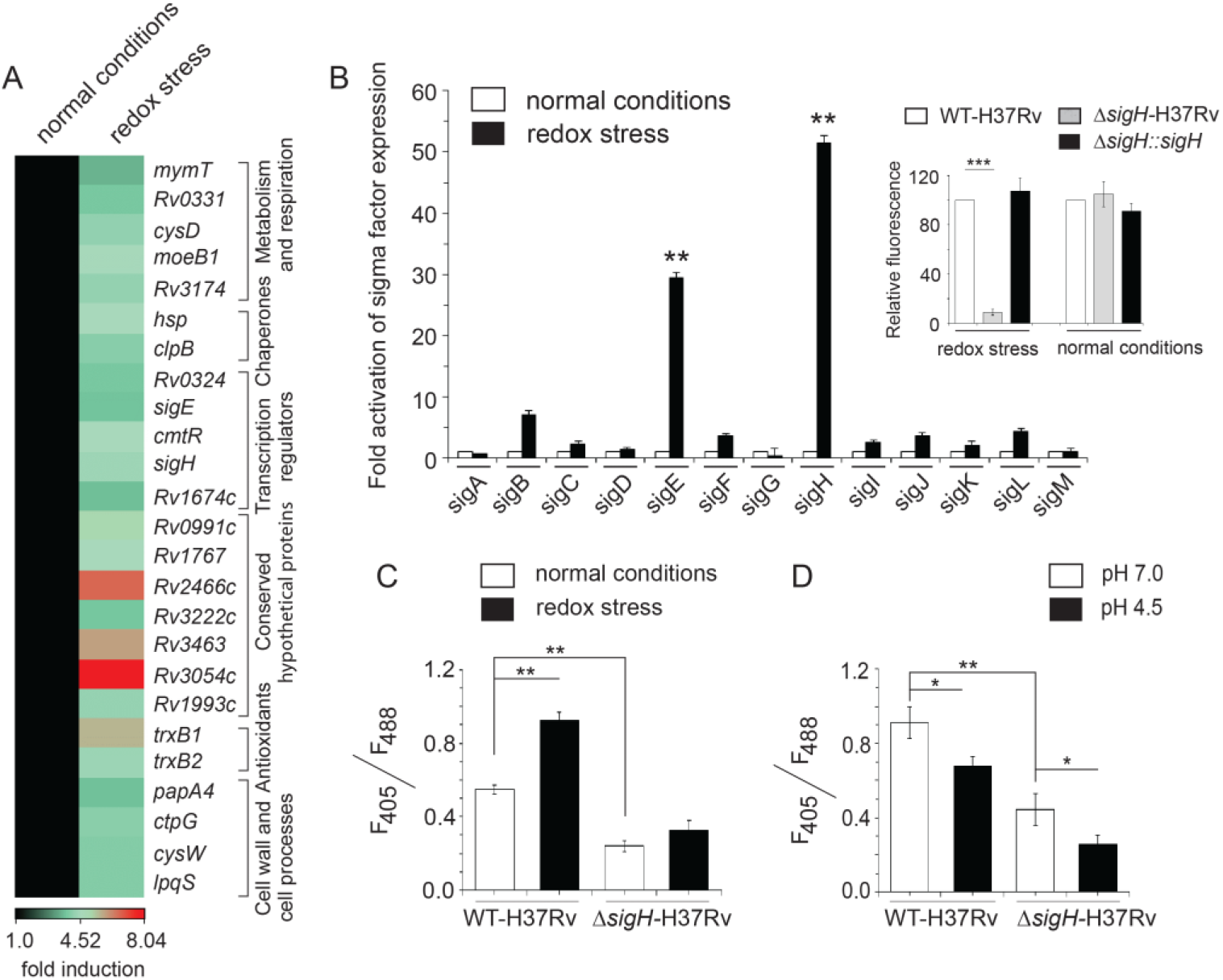
Redox stress-inducible mycobacterial genes. (A) RNA-sequencing data derived heat-map of WT-H37Rv treated with 5 mM diamide versus WT-H37Rv exposed to carry-over DMSO. The data shows a list of significantly induced genes (≥3.5-fold; p< 0.01), most of which belong to SigH regulon. (B) To investigate which sigma factors are induced in response to redox stress, expression of 13 mycobacterial sigma factor-encoding genes was compared by RT-qPCR using WT-H37Rv grown in absence or presence of 5 mM diamide (compare empty and filled columns) as described in the SI Methods. The results show average values from biological duplicates, each with two technical repeats (^**^P≤0.01). Non-significant differences are not indicated. Having shown that *sigH* displayed the highest level of redox-dependent activation, we compared growth of *ΔsigH* mutant with WT-H37Rv under redox stress (inset) using Alamar Blue assay (see SI Appendix, SI Methods). A significantly lower growth (<10%) of the mutant relative to the WT bacilli is consistent with the role of SigH as a major regulator of mycobacterial redox stress response. Note that bacterial survival under normal conditions of growth (in presence of carry-over concentrations of DMSO) and under redox stress (in presence of 2.5 mM diamide) were assessed by normalizing the data with respect to corresponding WT-H37Rv cultures (considered as 100%). (C-D) To examine if SigH contributes to mycothiol redox potential, we compared intra-mycobacterial mycothiol redox state of WT-H37Rv and Δ *sigH*-H37Rv grown under (C) normal conditions, versus redox stress, and (D) normal conditions (pH 7.0), versus acidic pH (pH 4.5). The biosensor response was measured by analysis of fluorescence emission at 510 nm after excitation of the samples at 405 nm and 488 nm. The data represent average values from biological triplicates, each with two technical repeats, suggesting that mycobacterial SigH regulates mycothiol redox potential (^*^P≤0.05; ^**^P≤0.01).

Having confirmed a striking activation of SigH under redox stress, we wanted to investigate impact of deletion of the sigma factor on susceptibility to redox stress. Thus, WT- and SigH-depleted H37Rv strains were grown in presence of increasing concentrations of diamide, and survival was monitored by using microplate-based Alamar Blue assay (SI Appendix, Fig. S2) as described above. To determine fold difference in susceptibility of WT and the mutant bacilli to diamide, we next quantified and plotted the fluorescence data (inset to Fig. 2B). Note that bacterial survival under normal conditions of growth (in presence of carry-over concentrations of DMSO) and under redox stress (in presence of 2.5 mM diamide) were assessed by normalizing the data with respect to corresponding WT-H37Rv cultures (considered as 100%). Importantly, *ΔsigH*-H37Rv showed a significant reduction in metabolic activity (> 10-fold) compared to the WT bacilli under redox stress in presence of 2.5 mM diamide. However, the mutant and WT bacilli grew comparably well under normal conditions of growth, and stable expression of SigH in the mutant bacteria could rescue redox-dependent growth defect. The above results prompted us to compare intra-mycobacterial mycothiol redox level in WT-H37Rv and *ΔsigH*-H37Rv. In keeping with the importance of *sigH* locus on mycobacterial survival under redox stress, the SigH-depleted mutant showed a significantly lowered mycothiol redox state relative to WT bacilli both under normal conditions and under redox stress (Fig. 2C). Furthermore, consistent with low pH-induced reductive stress and the importance of SigH under redox conditions of growth, Δ *sigH*-H37Rv displayed a lower mycothiol redox state (an increase in the reduced form of mycothiol) under acidic pH (Fig. 2D). These observations are in keeping with previous studies showing role of SigH in the induction of several genes involved in mycothiol redox homeostasis (41, 42). Taken together, these results suggest that both PhoP and SigH regulate physiological events leading to modulation of mycobacterial redox homeostasis.

### *phoP* controls global expression of genes associated with thiol redox homeostasis

In keeping with a significantly higher susceptibility of a PhoP-depleted mutant to redox stress (relative to the WT bacilli) (Fig. 1), we noted that a number of redox-inducible genes (Fig. 2A) are also regulated by the *phoP* locus (43). To identify the mechanism of functioning of PhoP, global transcriptional profiling was carried out by RNA sequencing using WT-H37Rv and *ΔphoP*-H37Rv, grown under redox stress. Fig. 3A shows the top 15 genes (>2-fold; p<0.05), which demonstrate highest level of activation in response to redox stress [SI Appendix, Table S3B (Excel spreadsheet)].

**Fig. 3:**
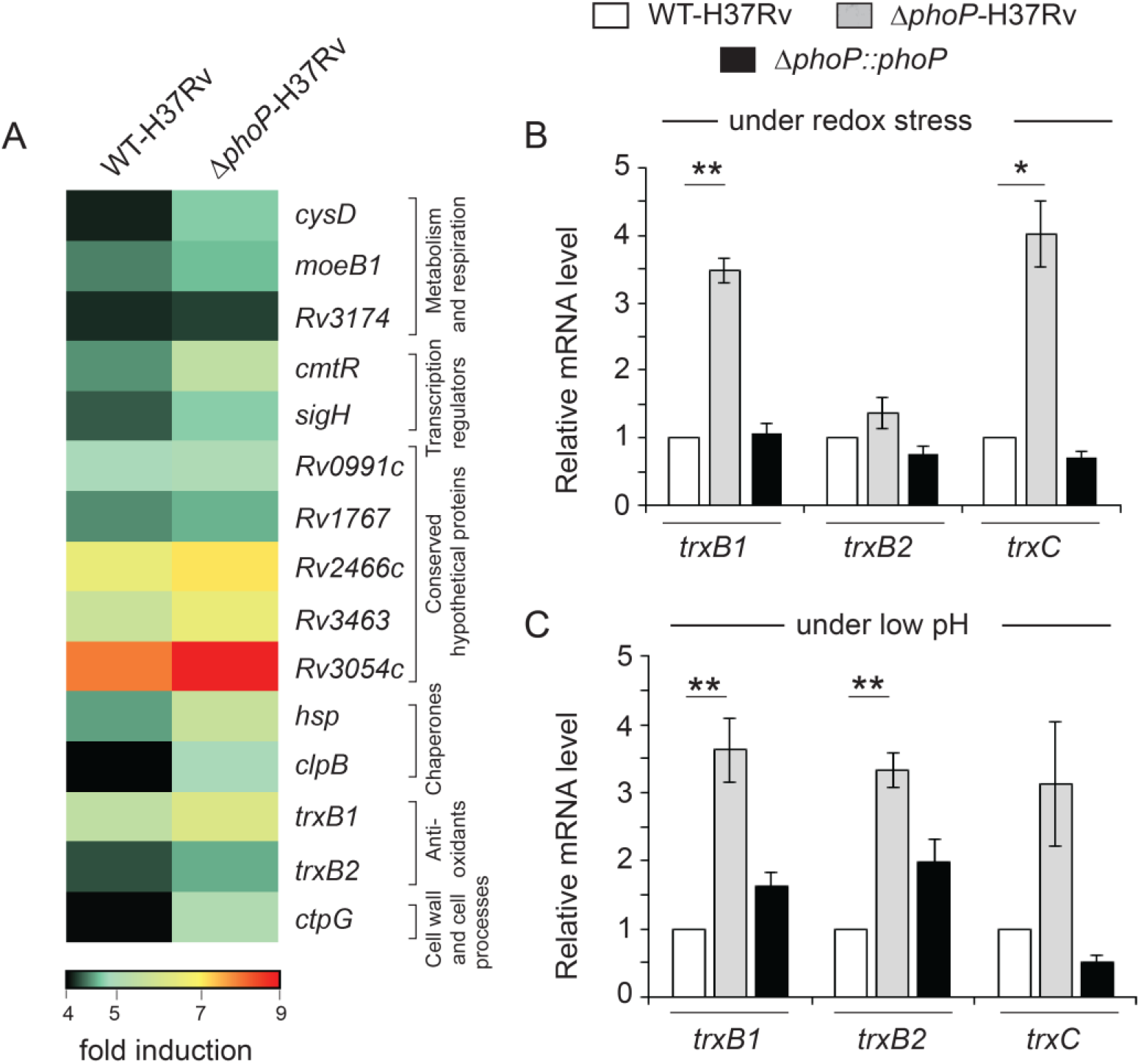
PhoP regulates expression of redox-inducible genes associated with SigH regulon. (A) RNA-sequencing derived heat-map of upregulated (≥2.0-fold; p< 0.05) genes in WT-H37Rv and Δ *phoP*-H37Rv, grown under under redox stress (as described in the SI Appendix) relative to their corresponding expression levels under normal conditions of growth (only top 15 genes are shown). The data shows a list of significantly induced genes, most of which belong to the SigH regulon (42). Notably, thioredoxin (*trx*) genes representing one of the major mycobacterial anti-oxidant systems are found to be part of the redox-active PhoP regulon. Although we observed a > 3-fold change in expression level of *trxC*, the differential expression was insignificant [SI Appendix, Table S3B (Excel spreadsheet)]. (B-C) Expression of redox-active thioredoxin genes in WT, *ΔphoP*-H37Rv, and complemented *ΔphoP*-H37Rv, grown either under redox stress (B) or under acidic pH conditions (C), were investigated by RT-qPCR measurements as described in the SI Methods. In all cases, fold differences in expression levels with standard deviations from replicate experiments were determined from at least three independent RNA preparations (^*^*P*<0.05; ^**^*P*<0.01).

Notably, we observed a significant overlap between PhoP-activated genes in presence of diamide and genes controlled by the alternative sigma factor SigH under redox stress (42, 44) or in presence of hydrogen peroxide (44). In keeping with the overlap between redox active PhoP and SigH regulon, numerous genes of SigH regulon, such as the thioredoxins (*trxB1, trxB2, trxC*), lipid synthesis genes (*papA4*), cysteine synthesis (*cysM*), and several predicted oxidoreductases (*Rv2454C, Rv3463*), showed a significant activation in Δ *phoP*-H37Rv relative to WT-H37Rv under redox stress [Fig 3A, and SI Appendix, Table S3B (Excel spreadsheet); also see Tables S3C-S3F]. These data strongly indicate that PhoP functions by targeting thiol homeostasis. Together, transcriptional profiling data suggest that PhoP plays a major role in thiol-associated stress response, and are in agreement with (a) disruption of redox-poise of *ΔphoP*-H37Rv under low pH as well as in presence of diamide and (b) enhanced sensitivity of *ΔphoP*-H37Rv to diamide.

Because redox-active thioredoxin genes belong to the PhoP-regulon (Fig. 3A), we next wanted to examine whether PhoP contributes to mycobacterial redox poise via its regulatory control of thioredoxin genes (*trxB1, trxB2* and *trxC*). Mycobacterial thioredoxin system, is one of the dedicated anti-oxidant systems to ensure growth and survival of the bacilli within the host (11, 12). Although bacterial thioredoxin reductases have been shown as drug targets (16, 45), the mechanism of regulation of thioredoxin gene expression remains largely unknown. Thus, we compared expression of mycobacterial thioredoxin genes in WT- and the PhoP-depleted H37Rv. In RT-qPCR experiments, *ΔphoP*-H37Rv showed a significantly higher expression of thioredoxin genes, particularly *trxB1* and *trxC* under redox conditions (Fig. 3B). More importantly, stable *phoP* expression in the complemented mutant could effectively restore expression of these genes to WT level. However, under normal conditions of growth, these genes in the mutant displayed a largely comparable expression as that of the WT bacilli (SI Appendix, Fig. S3). These results suggest that PhoP functions as a stress-specific repressor of *trx* genes, which control many important cellular processes. Since PhoPR is activated under acidic pH, a condition which causes reductive stress in mycobacteria (27, 29, 35), we next compared expression profile of thioredoxin genes in WT-H37Rv and *ΔphoP*-H37Rv, grown under low pH conditions (Fig. 3C). Importantly, under acidic pH conditions, thioredoxin promoters showed a significantly higher expression in *phoP*-H37Rv relative to WT bacilli. From these results, we conclude that PhoP is a stress-specific repressor of mycobacterial thioredoxin genes.

### Stress-specific SigH recruitment accounts for redox-dependent activation of thioredoxin promoters

Both PhoP-interacting extra-cytoplasmic sigma factors SigE and SigH (28) are significantly induced under redox stress (Fig. 2). Furthermore, consistent with previous reports our results suggest that SigH appears to regulate redox-active transcriptome of mycobacteria (40-42). In keeping with this, thioredoxin genes showed a significantly reduced expression in *ΔsigH*-H37Rv relative to the WT bacilli under redox stress and stable expression of SigH in the *sigH*-H37Rv could partly restore gene expression (Fig. 4A). Likewise, thioredoxin genes showed a reduced expression level in *ΔsigH*-H37Rv under low pH (Fig. 4B), but not under normal conditions of growth (Fig. 4C). Fig. 4D highlights the overlap of redox-inducible genes belonging to PhoP and SigH regulon, respectively [SI Appendix, Table S3B (Excel spreadsheet)]. As a control, expression of thioredoxin genes in WT-H37Rv and *ΔsigE*-H37Rv, grown under normal condition as well as under stress, reveal no significant difference in expression of *trx* genes (SI Appendix, Fig. S4).

**Fig. 4:**
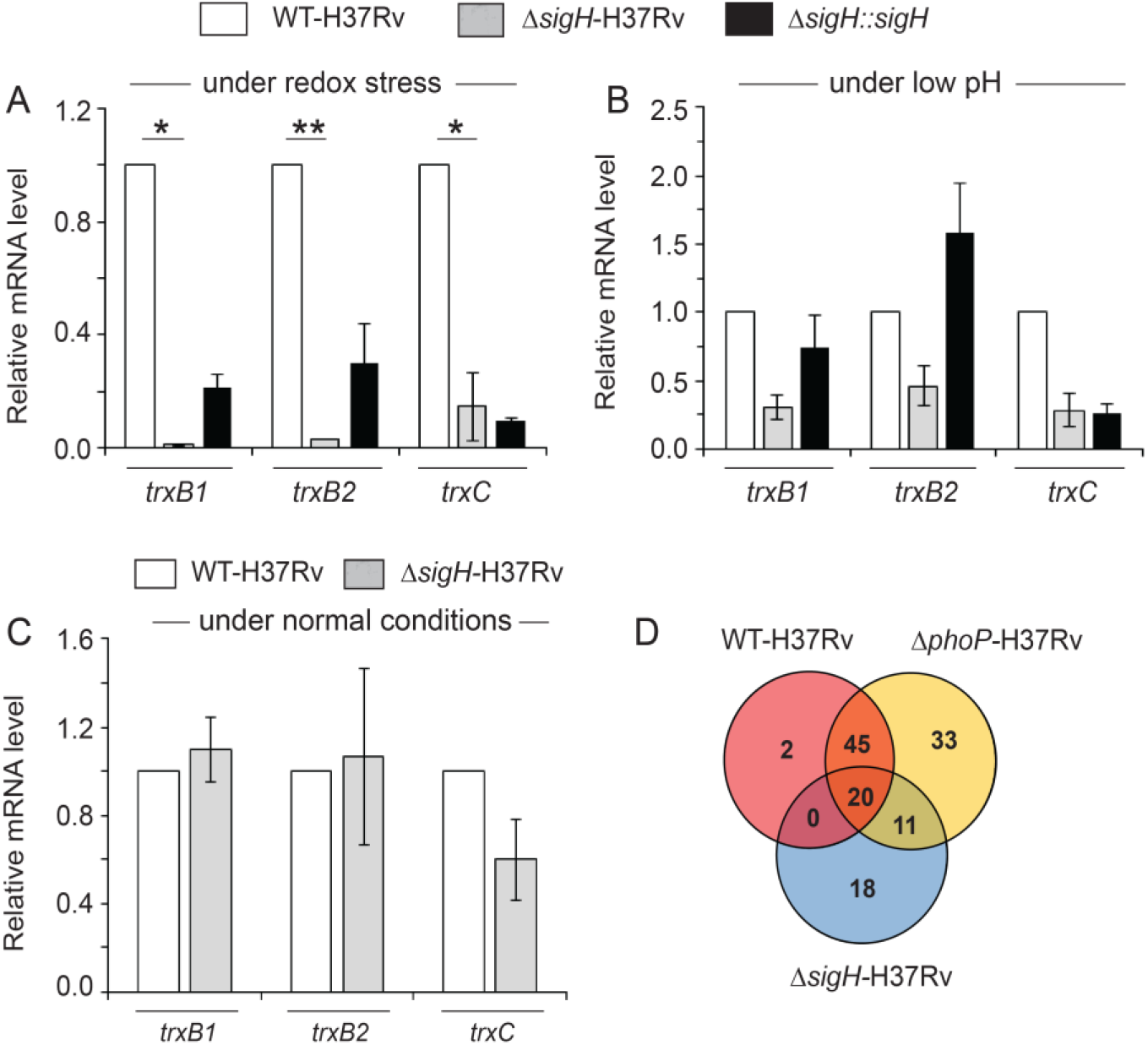
SigH functions as an activator of mycobacterial thioredoxin genes under stress. Expression of redox-active thioredoxin genes in WT, *ΔsigH*-H37Rv, and *ΔsigH::sigH*, grown under (A) redox stress, (B) under low pH and (C) under normal conditions of growth (as described in the SI methods) were examined by RT-qPCR (^*^*P*<0.05; ^**^*P*<0.01). The average fold difference in expression levels were determined from three independent RNA preparations as described in the SI Methods. (D) Venn diagram of genes upregulated (>2-fold; p< 0.05) in diamide treated WT-H37Rv and *phoP*-H37Rv (against their corresponding normal controls), as determined by RNA sequencing analysis [SI Appendix, Table S3C (Excel spreadsheet)], significantly overlap with the genes which belong to *sigH* regulon (42, 44). Data analyses involve comparison of genes annotated in the H37Rv genome alone, and the data represent average of two biological replicates.

Notably, PhoP-SigH interaction was unable to account for a contrasting regulation of thioredoxin gene expression, namely PhoP-dependent repression and SigH-dependent activation. While the thioredoxin promoters under redox stress were subdued in Δ *sigH* (Fig. 4A), these genes showed a noticeable induction in *phoP* (Fig. 3B). In agreement with the redox-dependent activation by SigH, also under low pH conditions we found a reproducible activation of thioredoxin genes by SigH (Fig. 4B). To investigate SigH recruitment within target promoters, we next expressed flag-tagged SigH in WT-H37Rv, and performed chromatin immunoprecipitation experiment (ChIP) followed by qPCR measurements (Fig. 5A). Our results demonstrate that under redox stress SigH is effectively recruited within thioredoxin promoters. However, no significant SigH recruitment was obtained within the identical promoters under normal conditions of growth (compare empty and filled columns, Fig. 5A). From these results, we conclude that stress-specific SigH recruitment within thioredoxin promoters maintains mycobacterial thiol redox homeostasis. In contrast, under identical experimental set up, we were unable to detect PhoP recruitment within the target promoters at either condition of mycobacterial growth (Fig. 5B). These results suggest that induction of redox-active thioredoxin promoter activity is attributable to direct recruitment of SigH within these promoters, and role of PhoP appears indirect in redox-dependent promoter activation. It should be noted that, ChIP data showing no significant recruitment of PhoP or SigH within these promoters under normal conditions, are consistent with a lack of regulatory expression of thioredoxin genes by these regulators under normal conditions of growth (SI Appendix, Fig. S3, and Fig. 4C).

**Fig. 5:**
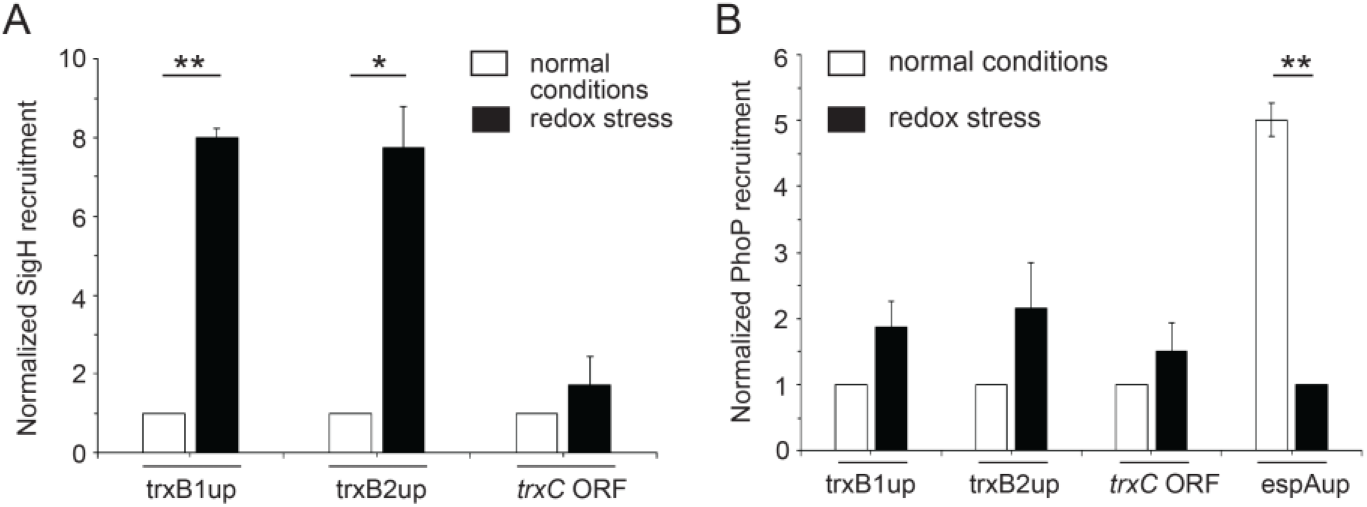
Redox-dependent recruitment of SigH within thioredoxin promoters. *In vivo* recruitment of (A) SigH and (B) PhoP within thioredoxin promoters was examined by ChIP-qPCR using WT-H37Rv grown under normal conditions and under redox stress. For ChIP assays, FLAG-tagged SigH or PhoP was expressed in WT-H37Rv as described in the SI Methods, and anti-FLAG antibody was used for immuno-precipitation. Note that trxC ORF refers to part of the *trxB2* gene, which remains adjacent to *trxC*, and espAup refers to the upstream regulatory region [-957 to -580 relative to the transcription start site of *espA* (Rv3616c)] comprising PhoP binding site (26). Fold PCR enrichment due to binding of the regulators to indicated promoters was determined with respect to an IP sample without adding anti-FLAG antibody (mock control). Each data was collected in duplicate qPCR measurements (technical repeats) using biological duplicates (^*^P≤0.05; ^**^P≤0.01). Note that each data was normalized relative to ChIP samples from cells grown under normal conditions. In contrast, as a positive control for examining PhoP recruitment by ChIP assays within espAup, IP samples from cells grown under redox stress was normalized to 1.

Importantly, PhoP-SigH interaction was unable to account for differential expression of thioredoxin genes. While expression of these genes under redox stress was noticeably activated in Δ *phoP*-H37Rv (Fig. 3B), under identical conditions these genes showed a clear repression in Δ *sigH*-H37Rv (Fig. 4A). To understand the mechanism of regulation, we sought to investigate whether any one of the two regulators control expression of the other. Thus, we compared *sigH* expression in *ΔphoP*-H37Rv, and *phoP* expression in *sigH*-H37Rv, respectively (SI Appendix, Figs. S5A, and Fig. S5C). In RT-qPCR experiments, under normal conditions SigH expression remains insignificantly higher in *ΔphoP*-H37Rv relative to WT-H37Rv (SI Appendix, Fig. S5A). In contrast, *phoP* expression is significantly induced in *ΔsigH*-H37Rv relative to WT-H37Rv, both under normal conditions and under redox stress [SI Appendix, Fig. S5C; Tables S3G and S3H (Excel spreadsheet)]. More importantly, stable SigH expression in *ΔsigH*-H37Rv complemented PhoP expression, suggesting that SigH functions as a repressor of PhoP expression. Together, these results suggest that stress-specific regulation of thioredoxin gene expression under redox stress is not attributable to PhoP-SigH interaction. In fact, these observations are consistent with expression profiles of thioredoxin genes in *ΔphoP*-H37Rv and *ΔsigH*-H37Rv under redox stress, displaying contrasting regulatory effects.

To investigate the physiological significance of PhoP-SigH interaction, we next designed an *ex vivo* experiment, where we studied SigH-dependent activation of *trxB1*, a representative promoter (also referred to as trxB1up). As expected, over-expression of SigH, in reporter assays using *M. smegmatis*, showed a significant promoter activation of *trxB1* relative to empty vector control (SI Appendix, Fig. S6). However, with expression of PhoP, we observed a significant inhibition of SigH-dependent activation of the promoter. As a control, *M. tuberculosis* RshA which functions as a negative regulator of SigH activity (46, 47), showed inhibition of SigH-dependent activation of trxB1up. However, expression of CFP-10 (as a negative control), under identical experimental set up, failed to inhibit SigH-mediated activation of trxB1up.

### PhoP interacts with redox-inducible mycobacterial sigma factors SigE and SigH

We previously showed that PhoP interacts with both SigE and SigH, the redox active sigma factors. While PhoP-SigE interactions have been implicated in pH homeostasis (28), the physiological consequence of PhoP-SigH interaction remains unknown. Since mycobacteria undergoes reductive stress under low pH conditions (35), we sought to further probe interactions between the extra-cytoplasmic sigma factors and PhoP. To this end, functional domains of PhoP were expressed in *M. smegmatis* along with SigE and SigH independently, and interaction between the mycobacterial regulators were examined by M-PFC (mycobacterial protein fragment complementation assay) (Fig. 6) as described previously (48). In this assay, the interacting proteins upon expression in *M. smegmatis* reconstitute functional DHFR (dihydrofolate reductase) and therefore, bacteria co-expressing interacting proteins can now grow on trimethoprim (TRIM) plates. Our M-PFC results show that while PhoPC interacts with SigH (Fig. 6A), it is the N-domain of PhoP (PhoPN) which interacts with SigE (Fig. 6B). To further validate the results, we performed *in vitro* pull-down assays (Fig. 6C-D). In this experiment, we have immobilized GST-tagged PhoP domains (PhoPN or PhoPC) on glutathione sepharose and incubated it with purified His-tagged SigH (Fig. 6C) or SigE proteins (Fig. 6D). GST-tag alone and glutathione sepharose beads, were used as controls. Importantly, results from pull down assays are in agreement with M-PFC data showing that N- and C-domains of PhoP interact with SigE and SigH, respectively. Together, these results suggest that the same molecule of PhoP possibly retains the ability to interact to both SigE and SigH simultaneously.

**Fig. 6:**
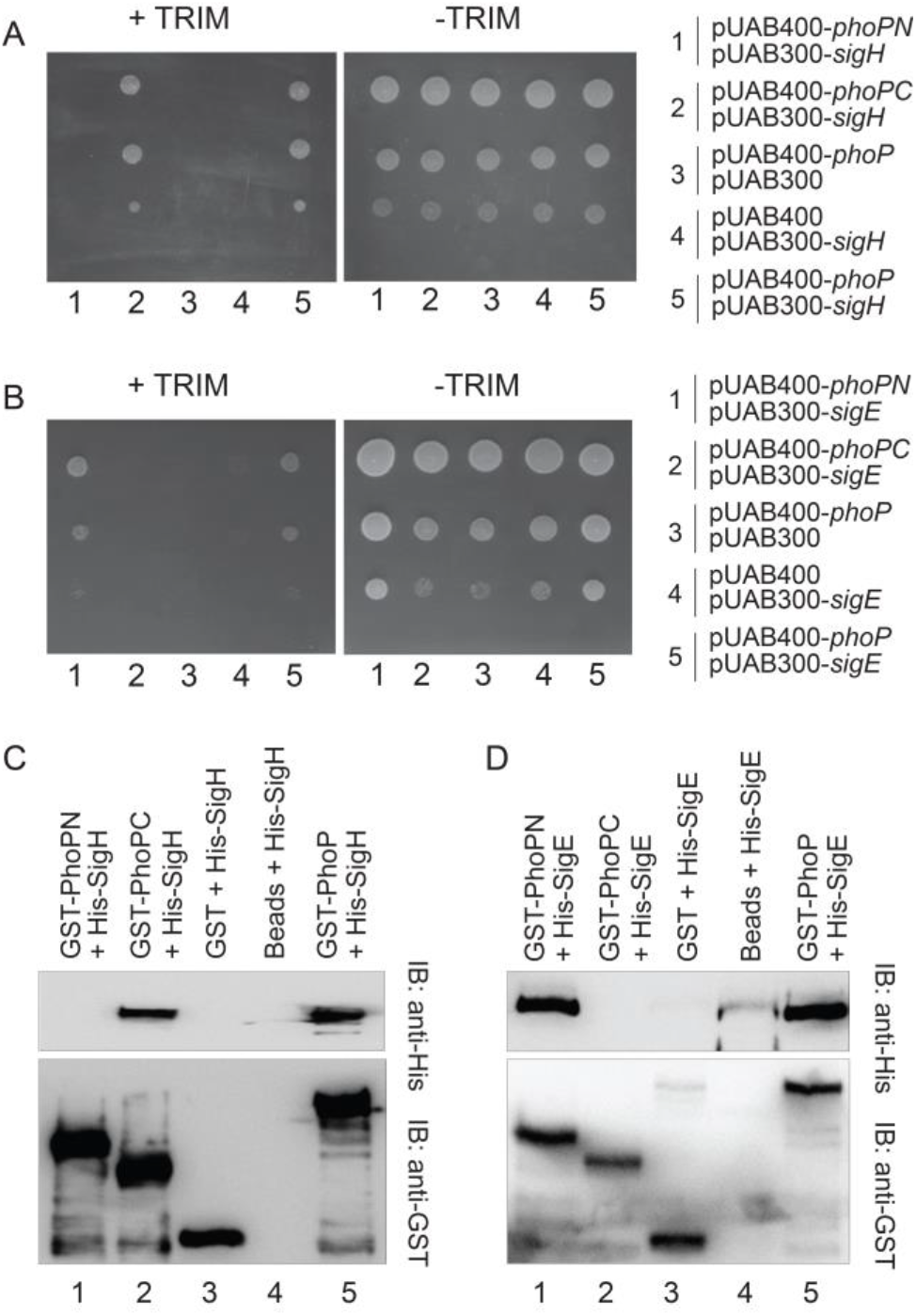
Probing interacting domains of PhoP-SigH. (A-B) In M-PFC experiments *M. smegmatis* co-expressing *M. tuberculosis* PhoP or its indicated domains and SigH (A) and SigE (B) were grown on 7H10/hyg/kan plates in the presence or absence of TRIM. Co-expression of fusions, pUAB400-*phoP*/pUAB300 and pUAB400/pUAB300-*sigH*, or pUAB400-*phoP*/pUAB300 and pUAB400/ pUAB300-*sigE* were used as respective empty vector controls. As positive controls, co-expression of pUAB400-*phoP*/pUAB300-*sigH* or pUAB400-*phoP*/pUAB300-*sigE* encoding PhoP/SigH or PhoP/SigE pairs, respectively, showed *M. smegmatis* growth in presence of TRIM. All of the strains grew well in absence of TRIM. To verify the above results by *in vitro* pull-down assays, recombinant His_6_-tagged SigH (C) or SigE (D) was incubated with glutathione-Sepharose, previously immobilized with GST-tagged PhoPN (lane 1), GST-tagged PhoPC (lane 2), GST alone (lane 3), resin alone (lane 4), or GST-tagged PhoP (lanes 5). Fractions of bound proteins (lane 1-5) were analysed by Western blot using anti-His (upper panel) or anti-GST antibody (lower panel). Together, these results suggest that PhoPN retains the ability to interact with SigE, whereas PhoPC interacts with SigH.

### Contrasting mode of stress-dependent PhoP-sigma factor interactions

We have shown above that SigH is recruited within thioredoxin promoters in a redox-dependent manner (Fig. 5A). However, under identical conditions recruitment of PhoP was undetectable (Fig. 5B). Mechanistically, this is in striking contrast to our previous results showing SigE recruitment within low pH -inducible promoters in a PhoP-dependent manner (28), where we showed recruitment of both PhoP and SigE within acid-inducible promoters during low-pH dependent mycobacterial adaptation. Therefore, it was of interest to investigate PhoP-SigH interaction *in vivo* under stress conditions. Thus, Δ *phoP*-H37Rv expressing a His-tagged PhoP, under the control of 19 kDa-antigen promoter in mycobacterial expression vector p19Kpro (49), was grown under normal and stress conditions (under low pH and in presence of 5 mM diamide, respectively). Next, whole cell lysates were incubated with Ni-NTA, and the bound protein was eluted as described previously (50). For cells grown under normal conditions, the eluent revealed a clear presence of SigH, suggesting *in vivo* interaction between PhoP and SigH (lane 1, Fig. 7A). This is in agreement with M-PFC results (Fig. 6A) and *in vitro* pull-down experiments (Fig. 6C) as described above. In striking contrast, we reproducibly noted almost undetectable signal of SigH in the eluent using cell lysates of identical strain, grown in presence of 5 mM diamide (lane 2, Fig. 7A). However, SigH was comparably present in mycobacterial cell lysates, grown under normal conditions and redox stress (lanes 3-4, Fig. 7A). Notably, SigE was comparably detectable in the eluent suggesting PhoP-SigE interactions under normal conditions as well as under redox stress (Fig.7A). Therefore, we surmise that PhoP interacts with SigE both under normal conditions and under redox stress. However, PhoP-SigH interaction occurs under normal conditions only, and is significantly reduced under redox stress.

**Fig. 7:**
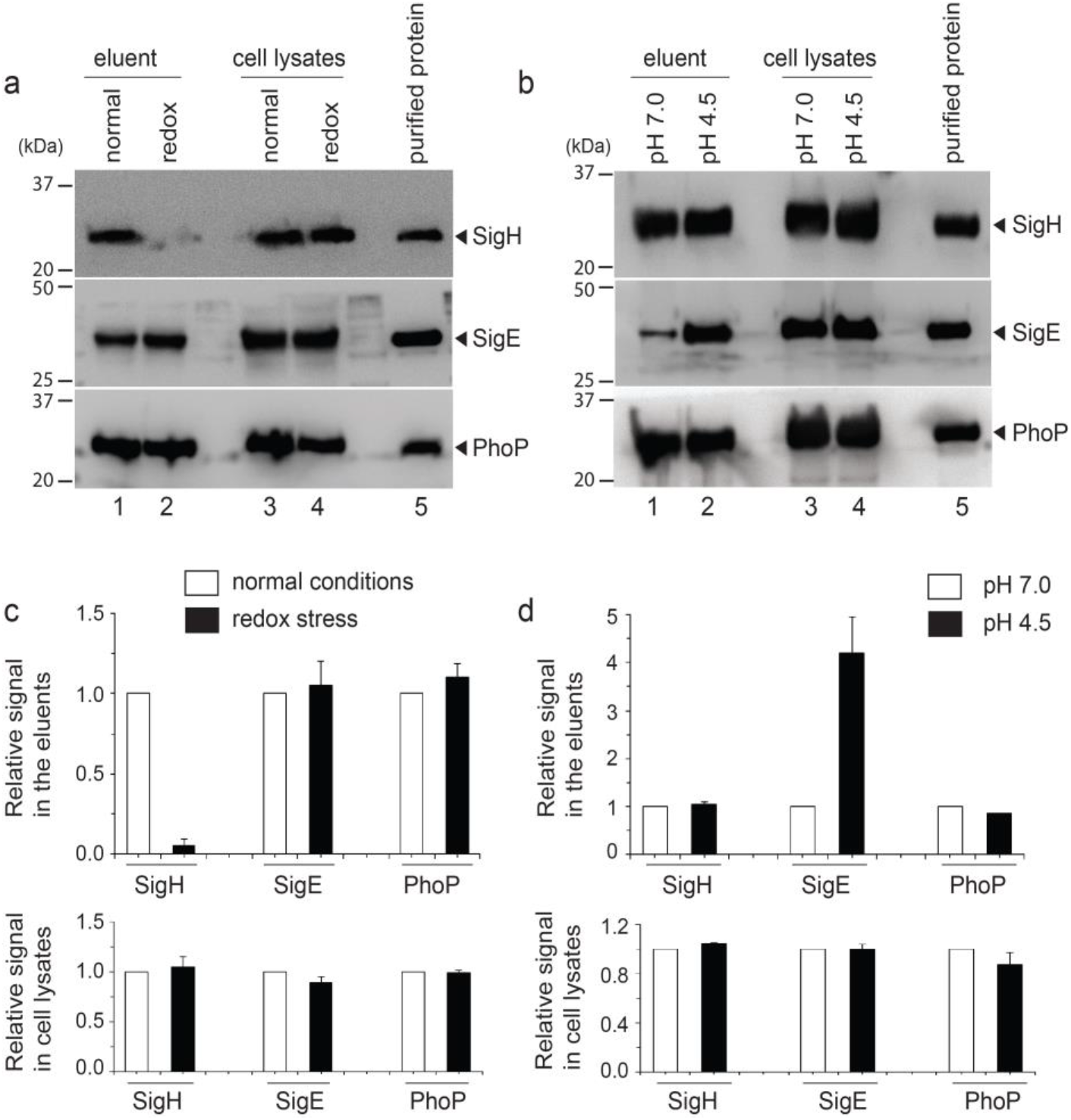
*In vivo* interactions of PhoP and extra-cytoplasmic sigma factors. To investigate stress-specific interactions of PhoP and sigma factors (SigE and SigH), *phoP*-H37Rv expressing a His-tagged PhoP, under the control of 19-kDa mycobacterial antigen promoter in p19Kpro (49), was grown under (A) normal conditions and redox stress (in presence of 5 mM diamide) or (B) normal conditions (pH 7.0) and acidic pH (pH 4.5), respectively. Next, the cell lysates were incubated with Ni-NTA, and bound protein was eluted. Panels A and B display immunoblots of eluents (lanes 1-2), corresponding cell lysates (lanes 3-4) and the purified protein (lane 5) detecting SigH, SigE and PhoP (from top to bottom), respectively in Western blots using antibodies directed against respective purified proteins. In each case, results are representative of two independent experiments. In some cases, the same blots were re-probed with multiple antibodies, signal intensity was quantified from replicate experiments and plotted below the gel images. Panels C and D refer to the plots derived from the data shown in panels A, and B, respectively.

Since mycobacteria under low pH encounters reductive stress, we next compared PhoP-SigH interactions in mycobacterial cells grown under normal and acidic pH (Fig. 7B). Importantly, under both conditions of growth, PhoP could interact with SigH. However, in agreement with our previous report (28), we observed a striking stimulation of PhoP-SigE interaction under acidic pH relative to normal conditions of growth. As expected, PhoP was present in the eluents as well as in the cell lysates, for cells grown under normal conditions, redox stress and low pH conditions, respectively. Figs. 7C-D show quantification of the data assessing stress-specific PhoP-SigH and PhoP-SigE interactions. These results are in agreement with two strikingly contrasting observations: (a) PhoP-independent redox-specific SigH recruitment within thioredoxin promoters (this study), and (b) PhoP-dependent low pH-inducible SigE recruitment within acid-inducible mycobacterial promoters of the PhoP regulon (this study and (28). Together, we conclude that stress-specific recruitment of extra-cytoplasmic sigma factors (SigE or SigH), controlled by PhoP-SigE/SigH interactions or lack thereof, mitigate stress response, regulate mycobacterial physiology under varying conditions of host environment, and enable intracellular survival of the bacilli.

## Discussion

*M. tuberculosis* activates expression of numerous proteins involved in redox homeostasis, including thioredoxins, alkyl hydroperoxide reductases, and the regulatory protein WhiB3. Importantly, PhoPR regulated *whiB3* expression (28, 51) controls synthesis of sulfolipids, poly- and diacyl trehaloses and phthiocerol dimycocerosate (PDIM) (52) and WhiB3-controlled lipid synthesis function as a reductive sink necessary to maintain redox homeostasis during acidic stress and macrophage infection (35). Thus, under low pH accumulation of mycobacterial NADH/NADPH generates redox stress, a metabolic requirement to oxidize these cofactors resulting in generation of reactive oxygen species (ROS). In line with these observations, *M. tuberculosis* under acidic pH display increased ROS synthesis, higher sensitivity to thiol stress, and a lower cytoplasmic redox potential relative to normal pH (35, 39). In another study, utilizing a redox-sensitive GFP-based probe to determine intra-mycobacterial thiol status, Abramovitch and co-workers showed that a reductive thiol environment was generated upon microbial growth using reduced carbon sources in the presence of acidic conditions (27). Notably, in this study a more reduced cytoplasm was reported for Δ *phoP*-H37Rv relative to WT bacilli. Further supporting an adaptive shift of redox poise under acidic pH, WT-H37Rv displayed a lower mycothiol redox state under acid stress (35). Together, these results suggest that under low pH, *M. tuberculosis* experience reductive stress, and a regulatory network of responses controlled by PhoPR and few other regulators mitigate reductive stress. However, the mechanism of coupling of pH stress with modulation of redox homeostasis remains obscure.

During its life cycle within the human host, *M. tuberculosis* encounters numerous stress conditions, such as hypoxia in animal models like mice, rabbit, guinea pig and nonhuman primates (53-55), increased levels of glutathione (GSH) inside lung granulomas of guinea pig (56), host generated nitric oxide (NO) in mice lung tissues and human tuberculous granulomas (54, 55) and carbon monoxide (CO) in mice lung and murine macrophages (57). Because of inhibition of respiration by NO or hypoxia (2), the NADH/NAD^+^ ratio goes up within the bacilli and *M. tuberculosis* encounters reductive stress (58). Consistent with this, metabolic profiling of granulomas from the lungs of infected guinea pigs suggests presence of increased GSH inside *M. tuberculosis*-laden granuolomas (56) and mycobacterial mutants defective in GSH uptake grow better within macrophages (59). These results suggest that despite tight regulation of host metabolic pathways, intracellular *M. tuberculosis* modulates its metabolic plasticity for survival and persistence (60). However, very little is understood about the underlying global regulatory mechanisms that integrates multiple environmental changes and intracellular gene expression of the bacilli.

In *M. tuberculosis* 10 out of 12 alternative sigma factors of a complex transcription machinery belong to extra-cytoplasmic function (ECF) family (40, 61), which control expression of genes in response to a variety of stress conditions (62, 63). We had uncovered that PhoP-SigE interaction accounts for mycobacterial pH homeostasis (28). Here, we explored the possibility of whether PhoP-SigH interaction contributes to mycobacterial redox homeostasis for the following reasons. First, coupling of pH stress and reductive stress has been shown to involve *phoPR* regulatory system; second, SigH has been implicated in regulating expression of thioredoxin genes, a major mycobacterial anti-oxidant system (41); third, global expression profiling of mycobacterial sigma factors under *in vitro* conditions of redox stress reveals a striking induction of SigH expression among all 13 sigma factors (Fig. 2). Along the line, previous studies have shown interactions between transcriptional regulators and sigma factors (64-66), contributing to regulation of numerous aspects of mycobacterial physiology.

We have previously shown that Δ *sigE*-H37Rv displayed a significantly lowered expression of acid inducible genes under conditions of low pH, suggesting a direct involvement of SigE in mycobacterial pH homeostasis (28). Here, we show that Δ *sigH*-H37Rv is significantly more susceptible to diamide (redox stress) relative to WT-H37Rv (inset to Fig. 2B). We extend these results to demonstrate that SigH is an activator of mycobacterial thioredoxin genes (Fig. 4A), and consistent with this, SigH is recruited within thioredoxin promoters in a redox-dependent manner (Fig. 5A), just as low pH-driven recruitment of SigE within acid-inducible *M. tuberculosis* promoters (28). However, despite interactions between PhoP and SigH (Fig. 6), SigH recruitment remains PhoP-independent (compare Fig. 5A and 5B). Upon further probing, we showed that *in vivo* PhoP and SigH interact under normal conditions, whereas the interaction is remarkably reduced under redox stress (Fig. 7A). Intriguingly, these results are not too surprising, since under redox stress SigH and PhoP display contrasting regulatory consequences of thioredoxin genes (compare Fig. 4 and Fig.3), a result we were unable to reconcile with PhoP-SigH interaction data.

PhoP functions as a repressor of *trx* genes (Fig. 3), whereas SigH regulates redox-specific activation of these genes (Fig. 4). Having found a contrasting regulatory effect, we probed whether any other control mechanism regulates expression of these promoters. Our finding that SigH appears to function as a repressor of PhoP expression [SI Appendix, Fig. S5C; Tables S3G and S3H (Excel spreadsheet] accounts for both SigH-dependent activation and PhoP-dependent repression of thioredoxin genes. The above results are now summarized in a schematic model (Fig. 8). According to this model, under normal conditions PhoP and SigH interact to each other. Therefore, possibly because of lower availability of free SigH, a functional SigH-bound RNAP is not formed. Thus, RNAP is unable to transcribe thioredoxin promoters effectively. Consistent with this notion, under normal conditions thioredoxin genes are poorly expressed, and there is no detectable SigH recruitment within thioredoxin promoters. In contrast, under redox stress SigH expression is strongly induced, and therefore, PhoP expression is significantly repressed. As a result, SigH-loaded RNAP transcribes thioredoxin genes and stress-specific SigH recruitment strongly induces expression of these genes to effectively mitigate redox stress. On the other hand, during low pH mycobacteria encounters reductive stress, thereby allowing SigE-loaded RNAP to transcribe PhoP-dependent acid-inducible genes. However, interaction between PhoP–SigH only involves a fraction of a free cellular pool of SigH, while a considerable part of free SigH remains available to mitigate reductive stress. The above model suggesting PhoP-dependent coordination of a major stress response, facilitates an integrated view of our results. Because PhoP shares different interaction interfaces with SigE and SigH (Fig. 6), our model further accommodates functioning of both the sigma factors under normal conditions of mycobacterial growth. Although our above results are consistent with a very high susceptibility of Δ *sigH*-H37Rv to redox stress (inset, Fig. 2B), given the fact that PhoP plays an indirect role in regulating expression of the major anti-oxidant system, we are unable to explain how a PhoP-depleted *M. tuberculosis* H37Rv is significantly sensitive to redox stress. We believe that many other PhoPR - regulated redox-active genes play a major regulatory role in mycobacterial redox stress response, and account for the mutant’s susceptibility to redox stress.

**Fig. 8:**
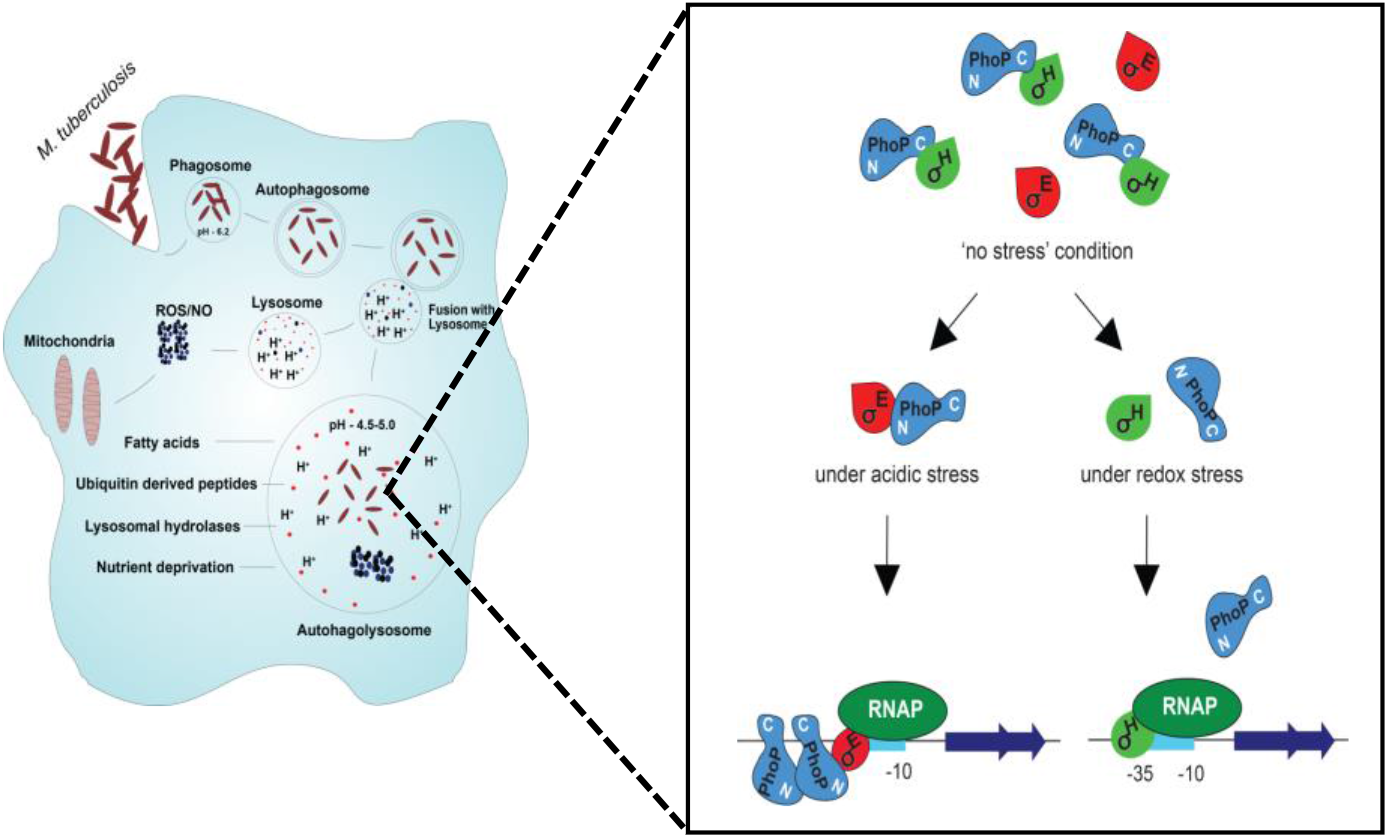
Schematic model depicting low pH- and redox-stress inducible mycobacterial gene expression. While PhoP restricts redox-inducible expression of mycobacterial thioredoxin genes, SigH is an activator of these genes under redox stress. Importantly, redox-inducible SigH expression appears to account for a significant repression of *phoP* expression. Thus, under redox stress PhoP-SigH interaction is of no physiological consequence and activation of thioredoxin genes is attributable to redox-dependent SigH recruitment. In agreement with this, under normal conditions thioredoxin genes are not influenced by either of the two regulators (PhoP or SigH). In contrast, under low pH conditions, both PhoP and SigE are recruited within acidic pH-inducible promoters of mycobacteria, and PhoP-SigE interaction within the transcription initiation complex promotes activation of acid-inducible genes. Consistently, unlike strikingly reduced interaction of PhoP-SigH under redox stress, we observed enhanced PhoP-SigE interaction under acidic conditions of growth. Taken together, PhoP-SigE interaction under acid stress contributes to pH homeostasis, whereas reduced PhoP-SigH interaction under redox stress, contributes to mycobacterial thiol redox homeostasis.

A large number of redox-inducible overlapping genes, regulated by both PhoP and SigH, include essential genes involved in metabolic pathways, respiration and expression of transcription regulators (Fig. 4D). These results indicate that most likely PhoP-SigH interactions is physiologically relevant when mycobacteria encounter redox stress. However, under normal conditions, interaction between the two regulators prevents unnecessary activation of SigH regulon. In fact, this is not too unexpected since activation of the SigH regulon consisting of ∼39 genes (42) would otherwise be energy demanding and therefore, a second molecular control mechanism (in addition to RshA-SigH) would be in place to precisely limit SigH activation, only under the most appropriate stress conditions.

While virulence regulator PhoP impacts expression of ∼2% of the *M. tuberculosis* genome, results reported here argue for a novel and unexpected mode of regulation on top of already complex regulatory mechanism, which distinguish regulators like PhoP from other members of the family. It would be worth investigating whether such sigma factor-dependent control of transcriptional regulators contributing to mycobacterial physiology remains a general feature involving several other sigma factors and regulators. Perhaps, these subtle features are related to the need for multiple regulatory functions during establishment and maintenance of infection. Given the nature of complex life-style of pathogenic mycobacteria, this may have evolved in order to further tune survival and growth of the pathogen in response to continuously changing complex physiology of the host.

## Materials and Methods

A full explanation of methods studied including bacterial strains and culture conditions, cloning of numerous constructs, expression and purification of proteins, mycothiol redox state estimation, RNA isolation, RNA-seq, library constructions and high throughput sequencing, RT-qPCR measurements, ChIP-qPCR studies, Alamar Blue assays, M-PFC assays, co-immunoprecipitation, immunoblotting, and β-galactosidase assays are detailed in the SI Appendix, SI Methods. The oligonucleotide primers used in RT-qPCR and ChIP experiments are listed in SI Appendix, Table S1. The primers used to generate recombinant constructs and plasmids used in this study are listed in SI Appendix, Table S2.

## Data Availability

All transcriptional profile files have been submitted to the GEO database at NCBI (accession number GSE171775).

## Supplemental data

The supplemental data include Supplemental Materials and methods, six supplemental figures, two supplemental tables, and one supplemental excel spreadsheet, which contains data for the Heat-map and Venn diagram showing genes that are differentially regulated (> 2-fold; p<0.05) either in WT-H37Rv or in the indicated mutants under normal conditions of growth or under specific stress conditions. Comparisons involve genes annotated in the H37Rv genome and data provided represent average value of two biological repeats.

## ACKNOWLEDGEMENTS

We thank Drs. G. Marcela Rodriguez and Issar Smith (The Public Health Research Institute, UMDNJ) for *ΔphoP*-H37Rv, complemented Δ*phoP*-H37Rv, Δ*sigE* and complemented Δ*sigE*-H37Rv strains, Riccardo Manganelli (University of Padova) for Δ*sigH*-H37Rv and complemented Δ*sigH* -H37Rv, Amit Singh (Indian Institute of Science) for Mrx1-roGFP2 plasmid, Adrie Steyn (University of Alabama) for pUAB300/pUAB400 plasmids, Prabhat Ranjan Singh for his help with the ChIP experiments and Vijjamarri Anil Kumar for preliminary results. Library construction, RNA-sequencing and data analysis have been carried out by AgriGenome Labs Private Limited (Cochin), India.

This work was supported by intramural funding from CSIR-IMTECH, and a research grant (to D.S) from SERB (EMR/2016/004904), Department of Science and Technology (DST). H.G., P.P., and H.K were supported by CSIR pre-doctoral fellowships. The funders had no role in the study design, data collection and interpretation, or the decision to submit the work for publication.

## Author contributions

H.G., and D.S. designed research; H.G., P.P., and H.K. performed research; H.G., and D.S. analyzed data; H.G., and D.S. conceived the idea; D.S. coordinated the whole study, and wrote the paper.

## References

1. J. A. Imlay, The molecular mechanisms and physiological consequences of oxidative stress: lessons from a model bacterium. Nat Rev Microbiol 11, 443–454 (2013).

2. A. Kumar et al., Redox homeostasis in mycobacteria: the key to tuberculosis control? Expert Rev Mol Med 13, e39 (2011).

3. V. H. Ng, J. S. Cox, A. O. Sousa, J. D. MacMicking, J. D. McKinney, Role of KatG catalase-peroxidase in mycobacterial pathogenesis: countering the phagocyte oxidative burst. Mol Microbiol 52, 1291–1302 (2004).

4. R. Bryk, C. D. Lima, H. Erdjument-Bromage, P. Tempst, C. Nathan, Metabolic enzymes of mycobacteria linked to antioxidant defense by a thioredoxin-like protein. Science 295, 1073–1077 (2002).

5. D. L. Piddington et al., Cu,Zn superoxide dismutase of Mycobacterium tuberculosis contributes to survival in activated macrophages that are generating an oxidative burst. Infect Immun 69, 4980–4987 (2001).

6. O. Dussurget et al., Role of Mycobacterium tuberculosis copper-zinc superoxide dismutase. Infect Immun 69, 529–533 (2001).

7. G. L. Newton et al., Distribution of thiols in microorganisms: mycothiol is a major thiol in most actinomycetes. J Bacteriol 178, 1990–1995 (1996).

8. V. Saini et al., Ergothioneine Maintains Redox and Bioenergetic Homeostasis Essential for Drug Susceptibility and Virulence of Mycobacterium tuberculosis. Cell Rep 14, 572–585 (2016).

9. T. Jaeger et al., Multiple thioredoxin-mediated routes to detoxify hydroperoxides in Mycobacterium tuberculosis. Arch Biochem Biophys 423, 182–191 (2004).

10. S. Nambi et al., The Oxidative Stress Network of Mycobacterium tuberculosis Reveals Coordination between Radical Detoxification Systems. Cell Host Microbe 17, 829–837 (2015).

11. O. Carmel-Harel, G. Storz, Roles of the glutathione- and thioredoxin-dependent reduction systems in the Escherichia coli and saccharomyces cerevisiae responses to oxidative stress. Annu Rev Microbiol 54, 439–461 (2000).

12. J. Lu, A. Holmgren, The thioredoxin antioxidant system. Free Radic Biol Med 66, 75–87 (2014).

13. C. Vilcheze et al., Mycothiol biosynthesis is essential for ethionamide susceptibility in Mycobacterium tuberculosis. Mol Microbiol 69, 1316–1329 (2008).

14. J. E. Griffin et al., High-resolution phenotypic profiling defines genes essential for mycobacterial growth and cholesterol catabolism. PLoS Pathog 7, e1002251 (2011).

15. Y. J. Zhang et al., Global assessment of genomic regions required for growth in Mycobacterium tuberculosis. PLoS Pathog 8, e1002946 (2012).

16. M. B. Harbut et al., Auranofin exerts broad-spectrum bactericidal activities by targeting thiol-redox homeostasis. Proc Natl Acad Sci U S A 112, 4453–4458 (2015).

17. K. Lin et al., Mycobacterium tuberculosis Thioredoxin Reductase Is Essential for Thiol Redox Homeostasis but Plays a Minor Role in Antioxidant Defense. PLoS Pathog 12, e1005675 (2016).

18. K. H. Rohde, R. B. Abramovitch, D. G. Russell, Mycobacterium tuberculosis invasion of macrophages: linking bacterial gene expression to environmental cues. Cell Host Microbe 2, 352–364 (2007).

19. E. Perez et al., An essential role for phoP in Mycobacterium tuberculosis virulence. Mol Microbiol 41, 179–187 (2001).

20. S. B. Walters et al., The Mycobacterium tuberculosis PhoPR two-component system regulates genes essential for virulence and complex lipid biosynthesis. Mol Microbiol 60, 312–330 (2006).

21. C. Martin et al., The live Mycobacterium tuberculosis phoP mutant strain is more attenuated than BCG and confers protective immunity against tuberculosis in mice and guinea pigs. Vaccine 24, 3408–3419 (2006).

22. J. Gonzalo Asensio et al., The virulence-associated two-component PhoP-PhoR system controls the biosynthesis of polyketide-derived lipids in Mycobacterium tuberculosis. J Biol Chem 281, 1313–1316 (2006).

23. R. Goyal et al., Phosphorylation of PhoP protein plays direct regulatory role in lipid biosynthesis of Mycobacterium tuberculosis. J Biol Chem 286, 45197–45208 (2011).

24. B. K. Johnson et al., The Carbonic Anhydrase Inhibitor Ethoxzolamide Inhibits the Mycobacterium tuberculosis PhoPR Regulon and Esx-1 Secretion and Attenuates Virulence. Antimicrob Agents Chemother 59, 4436–4445 (2015).

25. W. Frigui et al., Control of M. tuberculosis ESAT-6 secretion and specific T cell recognition by PhoP. PLoS Pathog 4, e33 (2008).

26. V. Anil Kumar et al., EspR-dependent ESAT-6 Protein Secretion of Mycobacterium tuberculosis Requires the Presence of Virulence Regulator PhoP. J Biol Chem 291, 19018–19030 (2016).

27. J. J. Baker, B. K. Johnson, R. B. Abramovitch, Slow growth of Mycobacterium tuberculosis at acidic pH is regulated by phoPR and host-associated carbon sources. Mol Microbiol 94, 56–69 (2014).

28. R. Bansal, V. Anil Kumar, R. R. Sevalkar, P. R. Singh, D. Sarkar, Mycobacterium tuberculosis virulence-regulator PhoP interacts with alternative sigma factor SigE during acid-stress response. Mol Microbiol 104, 400–411 (2017).

29. J. J. Baker, S. J. Dechow, R. B. Abramovitch, Acid Fasting: Modulation of Mycobacterium tuberculosis Metabolism at Acidic pH. Trends Microbiol 27, 942–953 (2019).

30. G. L. Newton et al., Characterization of Mycobacterium smegmatis mutants defective in 1-d-myo-inosityl-2-amino-2-deoxy-alpha-d-glucopyranoside and mycothiol biosynthesis. Biochem Biophys Res Commun 255, 239–244 (1999).

31. K. Van Laer et al., Mycoredoxin-1 is one of the missing links in the oxidative stress defence mechanism of Mycobacteria. Mol Microbiol 86, 787–804 (2012).

32. M. Rawat, C. Johnson, V. Cadiz, Y. Av-Gay, Comparative analysis of mutants in the mycothiol biosynthesis pathway in Mycobacterium smegmatis. Biochem Biophys Res Commun 363, 71–76 (2007).

33. M. Rawat et al., Mycothiol-deficient Mycobacterium smegmatis mutants are hypersensitive to alkylating agents, free radicals, and antibiotics. Antimicrob Agents Chemother 46, 3348–3355 (2002).

34. A. Bhaskar et al., Reengineering redox sensitive GFP to measure mycothiol redox potential of Mycobacterium tuberculosis during infection. PLoS Pathog 10, e1003902 (2014).

35. M. Mehta, R. S. Rajmani, A. Singh, Mycobacterium tuberculosis WhiB3 Responds to Vacuolar pH-induced Changes in Mycothiol Redox Potential to Modulate Phagosomal Maturation and Virulence. J Biol Chem 291, 2888–2903 (2016).

36. M. Mehta, A. Singh, Mycobacterium tuberculosis WhiB3 maintains redox homeostasis and survival in response to reactive oxygen and nitrogen species. Free Radic Biol Med 131, 50–58 (2019).

37. P. Tyagi, A. T. Dharmaraja, A. Bhaskar, H. Chakrapani, A. Singh, Mycobacterium tuberculosis has diminished capacity to counteract redox stress induced by elevated levels of endogenous superoxide. Free Radic Biol Med 84, 344–354 (2015).

38. M. Z. Khan et al., Protein kinase G confers survival advantage to Mycobacterium tuberculosis during latency-like conditions. J Biol Chem 292, 16093–16108 (2017).

39. G. B. Coulson et al., Targeting Mycobacterium tuberculosis Sensitivity to Thiol Stress at Acidic pH Kills the Bacterium and Potentiates Antibiotics. Cell Chem Biol 24, 993–1004 e1004 (2017).

40. S. Raman et al., The alternative sigma factor SigH regulates major components of oxidative and heat stress responses in Mycobacterium tuberculosis. J Bacteriol 183, 6119–6125 (2001).

41. J. D. Sharp et al., Comprehensive Definition of the SigH Regulon of Mycobacterium tuberculosis Reveals Transcriptional Control of Diverse Stress Responses. PLoS One 11, e0152145 (2016).

42. R. Manganelli et al., Role of the extracytoplasmic-function sigma factor sigma(H) in Mycobacterium tuberculosis global gene expression. Mol Microbiol 45, 365–374 (2002).

43. R. R. Sevalkar et al., Functioning of Mycobacterial Heat Shock Repressors Requires the Master Virulence Regulator PhoP. J Bacteriol 201 (2019).

44. M. I. Voskuil, I. L. Bartek, K. Visconti, G. K. Schoolnik, The response of mycobacterium tuberculosis to reactive oxygen and nitrogen species. Front Microbiol 2, 105 (2011).

45. J. Lu et al., Inhibition of bacterial thioredoxin reductase: an antibiotic mechanism targeting bacteria lacking glutathione. Faseb J 27, 1394–1403 (2013).

46. T. Song, S. L. Dove, K. H. Lee, R. N. Husson, RshA, an anti-sigma factor that regulates the activity of the mycobacterial stress response sigma factor SigH. Mol Microbiol 50, 949–959 (2003).

47. S. Kumar et al., Interaction of Mycobacterium tuberculosis RshA and SigH is mediated by salt bridges. PLoS One 7, e43676 (2012).

48. A. Singh, D. Mai, A. Kumar, A. J. Steyn, Dissecting virulence pathways of Mycobacterium tuberculosis through protein-protein association. Proc Natl Acad Sci U S A 103, 11346–11351 (2006).

49. K. A. De Smet, K. E. Kempsell, A. Gallagher, K. Duncan, D. B. Young, Alteration of a single amino acid residue reverses fosfomycin resistance of recombinant MurA from Mycobacterium tuberculosis. Microbiology 145 (Pt 11), 3177–3184 (1999).

50. P. R. Singh, A. K. Vijjamarri, D. Sarkar, Metabolic Switching of Mycobacterium tuberculosis during Hypoxia Is Controlled by the Virulence Regulator PhoP. J Bacteriol 202 (2020).

51. L. Feng, S. Chen, Y. Hu, PhoPR Positively Regulates whiB3 Expression in Response to Low pH in Pathogenic Mycobacteria. J Bacteriol 200 (2018).

52. A. Singh et al., Mycobacterium tuberculosis WhiB3 maintains redox homeostasis by regulating virulence lipid anabolism to modulate macrophage response. PLoS Pathog 5, e1000545 (2009).

53. H. P. Kuo et al., Nitric oxide modulates interleukin-1beta and tumor necrosis factor-alpha synthesis by alveolar macrophages in pulmonary tuberculosis. Am J Respir Crit Care Med 161, 192–199 (2000).

54. H. S. Choi, P. R. Rai, H. W. Chu, C. Cool, E. D. Chan, Analysis of nitric oxide synthase and nitrotyrosine expression in human pulmonary tuberculosis. Am J Respir Crit Care Med 166, 178–186 (2002).

55. M. I. Voskuil et al., Inhibition of respiration by nitric oxide induces a Mycobacterium tuberculosis dormancy program. J Exp Med 198, 705–713 (2003).

56. B. S. Somashekar et al., Metabolic profiling of lung granuloma in Mycobacterium tuberculosis infected guinea pigs: ex vivo 1H magic angle spinning NMR studies. J Proteome Res 10, 4186–4195 (2011).

57. A. Kumar et al., Heme oxygenase-1-derived carbon monoxide induces the Mycobacterium tuberculosis dormancy regulon. J Biol Chem 283, 18032–18039 (2008).

58. H. I. Boshoff et al., Biosynthesis and recycling of nicotinamide cofactors in mycobacterium tuberculosis. An essential role for NAD in nonreplicating bacilli. J Biol Chem 283, 19329–19341 (2008).

59. V. Venketaraman et al., Role of glutathione in macrophage control of mycobacteria. Infect Immun 71, 1864–1871 (2003).

60. A. Trivedi, N. Singh, S. A. Bhat, P. Gupta, A. Kumar, Redox biology of tuberculosis pathogenesis. Adv Microb Physiol 60, 263–324 (2012).

61. R. Manganelli, M. I. Voskuil, G. K. Schoolnik, I. Smith, The Mycobacterium tuberculosis ECF sigma factor sigmaE: role in global gene expression and survival in macrophages. Mol Microbiol 41, 423–437 (2001).

62. R. K. Karls, J. Guarner, D. N. McMurray, K. A. Birkness, F. D. Quinn, Examination of Mycobacterium tuberculosis sigma factor mutants using low-dose aerosol infection of guinea pigs suggests a role for SigC in pathogenesis. Microbiology 152, 1591–1600 (2006).

63. S. Rodrigue, R. Provvedi, P. E. Jacques, L. Gaudreau, R. Manganelli, The sigma factors of Mycobacterium tuberculosis. FEMS Microbiol Rev 30, 926–941 (2006).

64. Y. Hu, Z. Morichaud, A. S. Perumal, F. Roquet-Baneres, K. Brodolin, Mycobacterium RbpA cooperates with the stress-response sigmaB subunit of RNA polymerase in promoter DNA unwinding. Nucleic Acids Res 42, 10399–10408 (2014).

65. U. S. Gautam, K. Sikri, A. Vashist, V. Singh, J. S. Tyagi, Essentiality of DevR/DosR interaction with SigA for the dormancy survival program in Mycobacterium tuberculosis. J Bacteriol 196, 790–799 (2014).

66. J. Burian et al., The mycobacterial antibiotic resistance determinant WhiB7 acts as a transcriptional activator by binding the primary sigma factor SigA (RpoV). Nucleic Acids Res 41, 10062–10076 (2013).

